# Developmental mitochondrial Complex I activity determines lifespan

**DOI:** 10.1101/2023.06.21.545894

**Authors:** Rhoda Stefanatos, Fiona Robertson, Alejandro Huerta Uribe, Yizhou Yu, Kevin Myers, Beatriz Castejon-Vega, Tetsushi Kataura, L. Miguel Martins, Viktor I. Korolchuk, Oliver D.K. Maddocks, Alberto Sanz

## Abstract

Aberrant mitochondrial function has been associated with an increasingly large number of human disease states. Observations from *in vivo* models where mitochondrial function is altered suggest that adaptations to mitochondrial dysfunction may underpin disease pathology. We hypothesized that the severity of these maladaptations could be shaped by the plasticity of the system when mitochondrial dysfunction manifests. To investigate this, we have used inducible fly models of mitochondrial complex I (CI) dysfunction to reduce mitochondrial function at two stages of the fly lifecycle, from early development and adult eclosion. Here, we show that in early life (developmental) mitochondrial dysfunction results in severe reductions in survival and stress resistance in adulthood, while flies where mitochondrial function is perturbed from adulthood, are long-lived and stress resistant despite having up to an 75% reduction in CI activity. After excluding developmental defects as a cause, we went on to molecularly characterize these two populations of mitochondrially compromised flies, short- and long-lived. We find that our short-lived flies have unique transcriptomic and metabolomic responses which overlap significantly in discreet models of CI dysfunction. Our data demonstrate that early mitochondrial dysfunction via CI depletion elicits an adaptive response which severely reduces survival, while CI depletion from adulthood is not sufficient to reduce survival and stress resistance.

## Introduction

Mitochondria, the organelles that make multicellular life possible, regulate major cellular processes such as division, differentiation, and death (Nunnari and Suomalainen, 2012). The mitochondrion serves as the site of production for energy, signalling molecules and metabolites. At the core of this is oxidative phosphorylation, OXPHOS. OXPHOS requires a functional electron transport chain (ETC) made up of 4 multi-protein complexes (CI-CIV) to create a proton gradient that powers ATP synthase (CV). Complex I (CI) is the largest of the four ETC complexes. It not only acts as an electron entry point but a major source of NAD+ and mitochondrial ROS (mtROS). Deregulation and mutation of genes encoding CI subunits has been associated with a variety of disease states including diabetes, mitochondrial disease, cancer, and neurodegeneration (Pagliarini et al., 2008). It is therefore only right that CI is considered a highly relevant therapeutic target which regulates not only cellular but organismal fate (Stefanatos and Sanz, 2011).

The assembly, structure and function of CI have been intensively studied in several model systems. From these studies we have seen varied effects of deletion or depletion of CI on fitness, with the mechanisms underlying the consequences of dysregulated mitochondrial metabolism still unclear. In ageing, CI depletion has been shown to both reduce and extend lifespan in various model systems (Copeland et al., 2009b; Dillin et al., 2002; Grad and Lemire, 2004; Hegde et al., 2014; Kruse et al., 2008; Owusu-Ansah et al., 2013; Sanz et al., 2010). Currently, we do not fully understand how alterations of mitochondrial function can have such paradoxical effects. In the case of mitochondrial disease, primary mitochondrial CI deficiency can result in very severe disease that presents early and is often fatal, but it can also present mildly and later in life (Gorman et al., 2016; Gorman et al., 2015; Russell et al., 2020). The prevailing dogma proposes that mitochondrial dysfunction eventually results in an energy crisis that cannot be overcome. However, published works in distinct animal models have demonstrated that the effects of mitochondrial/CI dysfunction can be prevented or attenuated by interventions that do not directly target or rescue mitochondrial deficiency, and therefore energy production (Cerutti et al., 2014; Civiletto et al., 2018; Ferrari et al., 2017; Jain et al., 2019; Johnson et al., 2013). While characterising these models of mitochondrial dysfunction, a mountain of evidence has accumulated suggesting that the severity of pathology induced by mitochondrial/CI dysfunction depends on a factor unrelated to the dysfunction itself. We hypothesised that this “factor”, which determines the severity of the pathology is the mitochondrial plasticity of the organism, therefore developmental stage, when mitochondrial dysfunction manifests.

During development, organisms are highly adaptive to changes in the internal and external environment, with the mature organism carrying these adaptations into adulthood (Bazopoulou et al., 2019; Elkahlah et al., 2020; Gong et al., 2015; Gyllenhammer et al., 2020; Lafuente and Beldade, 2019). The *Drosophila* life cycle can be crudely divided into development and adulthood. Developing D*rosophila* have a greater capacity for regeneration and environmental adaptation than adult flies. To investigate whether inducing mitochondrial dysfunction at two distinct stages of the lifecycle would alter its effects, we created inducible fly models of mitochondrial complex I (CI) dysfunction. We decreased CI function during two periods of the fly life cycle, from early development or from eclosion (adults). Flies where mitochondrial dysfunction was induced developmentally had severely decreased lifespan and lower stress resistance, while flies where mitochondrial function was perturbed from adulthood were indistinguishable from control flies, despite a comparable decrease in CI activity. We were able to rule out developmental defects as a primary cause and using transcriptomic, metabolomic and proteomic approaches characterized these two populations of flies with compromised mitochondria. Short-lived mitochondrially compromised flies mount a unique biological response which overlaps significantly in discreet models of CI dysfunction but not with long-lived mitochondrially compromised flies. The data presented here demonstrate that the inherent plasticity of the organism when mitochondrial metabolism is challenged determines the grade of adaptive response which is mounted and therefore its effects on organismal fitness and lifespan.

## Materials and Methods

### Fly husbandry

Flies were maintained on standard media (1% agar, 1.5% sucrose, 3% glucose, 3.5% dried yeast, 1.5% maize, 1% wheat, 1% soya, 3% treacle, 0.5% propionic acid, 0.1% Nipagin) at 25°C in a controlled 12Lh light: dark cycle. Male flies were used in all experiments unless otherwise stated. For *GeneSwitch* experiments flies were placed on media supplemented with either vehicle (EtOH) or varying concentrations of RU486 (Abcam, ab120356). In experiments where the Gal4/Gal80^ts^ system was utilised, flies were maintained at either 18 or 29°C.

Experimental flies were collected within 48 hrs of eclosion using CO_2_ anaesthesia and kept at a density of 20 flies per vial. Genotype and source information for fly lines used in this study can be found in supplementary Table 1.

### RU486 feeding protocol

To activate *tubGS* from development, mating adults were flipped onto media containing titrated concentrations of RU486 (up to 0.5µM). To activate *tubGS* from adulthood, male progeny were collected and placed on standard media containing 500µM RU486.

### Survival analysis

Survival graphs were created using GraphPad Prism 9 with log-rank statistics applied. A minimum of 50 flies were used for each experimental group and two-three replicate experiments were performed. Graphs presented depict pooled/representative results.

#### Lifespan 25°C and 29°C

Flies were collected within 24 hours of eclosion, maintained at a density of 20 flies per vial at the appropriate temperature (25°C or 29°C). They were transferred to fresh media every 2-3 days and the number of dead flies was scored.

#### Hydrogen peroxide treatment

3-5 day old adult male flies were starved overnight and then transferred to vials containing 5ml 1% agar, 5% Sucrose, 2.5% H_2_0_2_. The number of dead flies were scored at regular intervals.

#### Thermal Stress

Flies were collected within 24 hours of eclosion, maintained at a density of 20 flies per vial at 32°C. They were transferred to fresh media every 2-3 days, and the number of dead flies was scored.

#### Starvation

To assay starvation, 3-5 day old adult male flies were placed into vials containing 2ml of 1% agar. The number of dead flies was scored at regular intervals.

### CI-linked respiration

Mitochondrial oxygen consumption was measured in homogenates of whole male L3 larvae and 3–5 day old adults using a Oxygraph 2-K (Oroboros Instruments). 5 L3 larvae/10 flies were homogenised in isolation buffer (250 mM sucrose, 5 mM Tris–HCl, mM EGTA, pH 7.4) and diluted 10x in assay buffer (120 mM KCl, 5 mM KH2PO4, 3 mM HEPES, 1 mM EGTA, 1 mM MgCl2, 0.2% (w.v.) BSA, pH 7.2). For L3 larvae the following substrates were added, 5mM Proline, 5mM Pyruvate, 2.5mM Glutamate, 2mM Malate. For adults 5mM Proline, 5mM Pyruvate was added. State 3 was initiated with the addition of 1mM ADP. Raw O_2_ Flux extracted using Oroboros DatLab 7.0 was normalised to the protein concentration (quantified via Bradford assay) of each homogenate (picomoles of O_2_ per min^-1^ per mg^-1^).

### Western blot

3-5 day old males of the indicated genotypes and experimental groups were homogenised in 150µl of homogenisation buffer (0.2%Triton X-100 supplemented with 1 x complete mini EDTA-free protease inhibitor (Roche) in PBS) using a motorised pestle homogeniser for ∼1min. After a 10min incubation at 4°C, samples were centrifuged at 13,000 rcf for 15 mins at 4°C. Protein concentration of the supernatants were quantified via the Bradford (ThermoFisher) assay and a FLUOstar Omega plate reader (BMG Labtech). Samples were mixed 1:1 with 2x Laemmli sample buffer (BioRad) supplemented with 5% □-ME (Sigma) and boiled at 95 °C for 5min. 30µg of protein was run on Any kD™ Criterion™ TGX Stain-Free™ Protein Gels (BioRad) and transferred to nitrocellulose membranes (Amersham™ Protran®) using the Criterion™ Blotter wet transfer system (BioRad). Membranes were blocked in 5% Milk in PBS, 0.1% Tween-20 for 1 hour at room temp. They were then washed with PBS and incubated with 1° antibodies (α-NDUFV2, 15301-1-AP, Protein tech, α-tubulin, Abcam, ab179513) in 5% milk 0.1% Tween 20, overnight at 4°C. Membranes were washed 3x for 10 mins with PBS with 0.1% Tween 20. This was followed by incubation with HRP conjugated 2° antibodies (α-mouse (PI-2000) and α-rabbit (PI-1000, Vector Labs)) in 5% Milk in PBS, 0.1% Tween-20 for 1hr at room temp. Membranes were washed as above. Hybridisation was visualised via Clarity Western ECL Substrate (BioRad) and using a LAS-4000 CCD camera system.

### qPCR gene expression analysis

Total RNA was extracted in quadruplicate from male L3 larvae and 3-5 day old adults of the indicated genotypes and experimental groups using standard Trizol (ThermoFisher) extraction. RNA was DNAse I (ThermoFisher) treated and purified via ethanol precipitation. cDNA was synthesised using the High-Capacity cDNA reverse transcription kit (Applied Biosystems) according to manufacturer’s instructions. Relative expression of the target genes was quantified using QuantiNova SYBR green reagents and Applied Biosystems Step One qPCR instrument. Data were extracted and analysed using Applied Biosystems Step One software version 2.3 and GraphPad Prism 9. Relative expression of the target genes (*nd-18* and *nd-75*) was normalised to levels of the house keeping gene *act88f*. For each target, mean fold change with standard error mean is presented. Primer sequences can be found in supplementary Table 2.

### RNA sequencing and differential expression analysis

For RNAseq, total RNA was extracted as outlined in qPCR gene expression analysis. RNA samples were submitted to the University of Glasgow Polyomics facility for QC (Agilent), PolyA selection, Library preparation (TruSeq Stranded mRNA Library Prep) and sequencing using the Illumina Next Seq platform (NextSeq 2000 P3 Reagents, NextSeq 500/550 High Output Kit v2.5). Upon receipt of raw data, reads were trimmed mapped and counted using an in-house pipeline (described in supplementary table 3, scripts available upon request). Differential expression analysis was performed by using a custom R script (available upon request) utilising the limma package (Ritchie et al., 2015).

For GeneOntology and Pathway Analysis, Gene names were retrieved from Flybase, before being sorted by FDR adjusted p-value (q-value). Genes were determined to be significant if they had a q-value less than 0.05. The significant gene names, ranked by their q-value, were then submitted to FlyEnrichr (Chen et al., 2013) where the results for GO Biological Process, Cell Component, Molecular Function 2018 and KEGG 2019 were assessed and ranked by Combined Score to establish the most representative pathways in each comparison.

All graphics associated with the DE analysis were created in R Studio, a list of all modules used within their creation can be found in Supplementary Table S3. The R script for the creation of the graphics is also available on request.

### Metabolomics

Metabolomic analysis via LC-MS was performed on quadruplicate samples of 3-5 day old male adults of the indicated genotypes and experimental groups. Anaesthetised flies were homogenised in ice cold extraction buffer (50% HPLC grade methanol (Fisher), 30% HPLC grade acetonitrile (Sigma), 20% water) and centrifuged at 15,000 rpm at 4°C. Supernatants were stored at −80C and transferred to LC-MS vials before analysis. LC-MS and data analysis was performed as previously described (Maddocks et al., 2017). Analysis of peak data was performed using MetaboAnalyst 5.0(Pang et al., 2021).

### Proteomics

Mass spectrometry analysis was performed at the Proteomics Facility of the Medical Research Council Toxicology Unit University of Cambridge, Cambridge, UK. Briefly, 50 5-day old adult males were homogenised in 450µL of 100mM Triethylammonium bicarbonate (TEAB) for 5 cycles of 20s at 5500rpm with 10s interval at 4°C. 1% RapiGest buffer and a further 450µL of 100mM TEAB buffer was added to the samples followed by sonication and centrifugation at 2000g for 5min at 4°C. 850µL of supernatant was collected for further processing. 50µg of protein extract was digested in 50µL 100mM TEAB. Samples were denatured at 80°C for 10min. 2.8μl of 72mM DTT was added, and samples were heated at 60°C for 10 min. 2.7μl of 266 mM Iodoacetamide was added and samples were incubated at room temperature for 30 minutes. DTT concentration was raised to 7mM to quench alkylation. Samples were digested with 1µg of trypsin (5µL of 0.2µg/µL stock) overnight at 37°C. Digests were quantified using the Peptide quantification assay from Pierce™ Quantitative Colorimetric Peptide Assay (Cat no: 23275) according to manufacturer instructions. 2xTMT11 with pooled reference standards were used to multiplex samples. LC-MS/MS using synchronous precursor selection (SPS MS3) with MS3 based quantification triggered by Drosophila MS2 identified peptides real time search was performed. Injected samples were analysed using an Ultimate 3000 RSLC™ nano system (ThermoFisher, Hemel Hempstead) coupled to an Orbitrap Eclipse™ mass spectrometer (ThermoFisher). Data was acquired using three FAIMS CVs (−45v, −60v and −75v) and each FAIMS experiment had a maximum cycle time of 2s. For each FAIMS experiment the data-dependent SPSMS3 RTS program used for data acquisition consisted of a 120,000 resolution full-scan MS scan (AGC set to 50% (2e5 ions) with a maximum fill time of 30ms) using a mass range of 415-1500 m/z. Raw data were imported and processed in Proteome Discoverer v2.5 (ThermoFisher). The raw files were submitted to a database search using Proteome Discoverer with Sequest HT against the Uniprot UP000000803 *Drosophila* database (containing 21135 seq. accessed on 2022/08/04). Common contaminant proteins (human keratins, BSA and porcine trypsin) were added to the database. Statistical analyses for fold change and significance were performed similarly as a previously described (Popovic et al., 2021).

### Immunofluorescence and Confocal microscopy

Adult fat bodies were dissected and fixed in 4% paraformaldehyde at room temperature for 30 mins. Tissues were first rinsed in 1xPBS and then incubated with LipidTOXred 1:500 and DAPI (1:1000) in 1x PBS + 0.005% Saponin for 1hr at room temp. Tissues were then washed 3x for 15 mins in 1x PBS + 0.005% Saponin and mounted with Prolong Diamond on polylysine coated glass slides with 13mm x 0.12mm spacers. Confocal images were taken on a Leica SP8 DLS and processed using Image J.

### TAG assay

4 x 15 males adult flies per indicated genotype and condition were weighed and then homogenised in Tet buffer (10 mM Tris, 1 mM EDTA, pH 8.0 with 0.1% v/v Triton-X-100) using glass beads and an OMNI Bead Ruptor. Samples were centrifuged at 13000 rcf for 5 min at 4°C. The supernatant was incubated at 72°C for 15mins. 3µl of sample and glycerol standards was added to 300µl of ThermoFisher Infinity Triglyceride reagent and incubated for 15m at 37°C. Absorbance at 540nm was measured on a Multiskan FC (ThermoFisher). A glycerol standard curve, weight and number of flies was used to calculate TAG/mg/fly.

### Transmission Electron Microscope

Thoraces from 5 male 3-5 day old flies of indicated genotypes and conditions were dissected in ice cold PBS and put directly into glutaraldehyde fixing solution O/N. Sample processing and preparation of sections was carried out by the Newcastle University EM facility. Ultrathin longitudinal sections of indirect flight muscle were analysed by TEM using a Hitachi HT7800 120kV TEM with the help of NU EM facility staff and processed using Image J.

### Histology

Fly heads were dissected and fixed overnight in 10% formalin. Heads were mounted in between two layers of 1% agar. These were further fixed in 10% formalin overnight and then paraffin embedded at the Newcastle University Biobank. Paraffin blocks were trimmed until the fly head became visible. 4µm sections were mounted onto polylysine slides and processed for heamotoxylin and eosin staining by the Newcastle University Biobank. Imaging was done using a Leica DMi8 and processed using Image J.

### NAD/NADH measurements

Measurements of NAD+ and NADH in whole flies were performed as described in (Kanamori et al., 2018; Kataura et al., 2022). Briefly, NAD+ or NADH were extracted from triplicates of 10 (NAD+) or 20 flies (NADH) homogenised with a motorised pestle using 20% trichloroacetic acid (TCA) (Sigma) or 0.5 M sodium hydroxide (NaOH, Sigma) 5 mM EDTA (Sigma) respectively. Sample pH was adjusted to 8.0 with 1M Tris. Levels of NAD+ and NADH were determined by levels of resorufin fluorescence produced from the enzymatic cycling reaction using resazurin (Sigma), riboflavin 5’-monophosphate (Sigma), alcohol dehydrogenase (Sigma) and diaphorase (Sigma). Fluorescence intensity was measured once a minute for a total of 60 minutes with a FLUOstar Omega, (BMG Labtech). NAD+ and NADH levels were determined by a β-NAD (Sigma) standard curve and adjusted to protein concentration determined by the DC protein assay (BioRad).

### Statistical Analysis

For all bar graphs and survival curves were produced using GraphPad Prism 9 with students *t*-test and Log-Rank test applied where appropriate. All data presented is the mean with SEM.

### Schematic Diagrams

All schematic diagrams were created using the biorender.com platform.

## Results and Discussion

### Ubiquitous developmental but not adulthood depletion of Complex I shortens lifespan

To understand whether adaptative plasticity could be determining the severity of pathology associated with mitochondrial dysfunction, we employed novel inducible fly models of mitochondrial dysfunction.

Using a combination of the GeneSwitch expression system (Roman et al., 2001) and RNAi interference against subunits of CI, we induced whole body mitochondrial dysfunction from either development (D+A) or adult eclosion (A only). This allowed us to directly test the effects of mitochondrial dysfunction when adaptative plasticity is high, i.e., development and when it is low, i.e., adulthood. Experimental flies were derived from crossing a ubiquitous GeneSwitch driver *tubulinGS* with either a control line (UAS-Empty/Ctrl) or RNAi lines targeting two different subunits of CI, ND-18 (ND-18 KD) and ND-75 (ND-75 KD). ND-18, orthologue of *Ndufs4*, is an accessory subunit of CI required for assembly and stability (Garcia et al., 2017). ND-75, orthologue of *Ndufs1*, is an essential subunit required for assembly, stability, catalytic activity, and super complex formation (Garcia et al., 2017). Both subunits have been found to be altered in patients suffering from mitochondrial disease(Benit et al., 2001; van den Heuvel et al., 1998).

To induce knockdown from development (D+A), flies were crossed on fly media containing appropriate concentrations (drug food) of the inducer (RU486) and upon eclosion maintained in vials containing drug food (detailed in material and methods) (Figure 1A). To limit knockdown to the adult phase (A only), flies were crossed on normal fly media and transferred to drug food after eclosion (Figure 1A). We confirmed the depletion of ND-18 and ND-75 at the mRNA and CI at the protein level in adult male flies (Figure S1A and C) and late-stage male larvae (L3) (Figure S1B). We saw no difference in the levels of depletion between D+A and A only ND-18 KD, ND-75 KD, or Ctrl flies (Figure S1A, B, C and D). The D+A group in both ND-18 and ND-75 KD flies were severely short-lived in comparison to A only KD flies, as well as D+A and A only Ctrl flies (Figure 1B, C and S1E compare solid line (A only) with dashed line (D+A)). These stark differences in survival were also observed in female D+A vs A only ND-18 KD and ND-75 KD flies (Figure S1F and G). This is particularly interesting given conflicting published reports where manipulation of CI has both reduced and extended lifespan (Cho et al., 2011; Copeland et al., 2009a; Scialo et al., 2016). Our data, reconcile previously contradictory findings, and indicate that fully functional CI is necessary during development but not adulthood.

**Figure 1.**
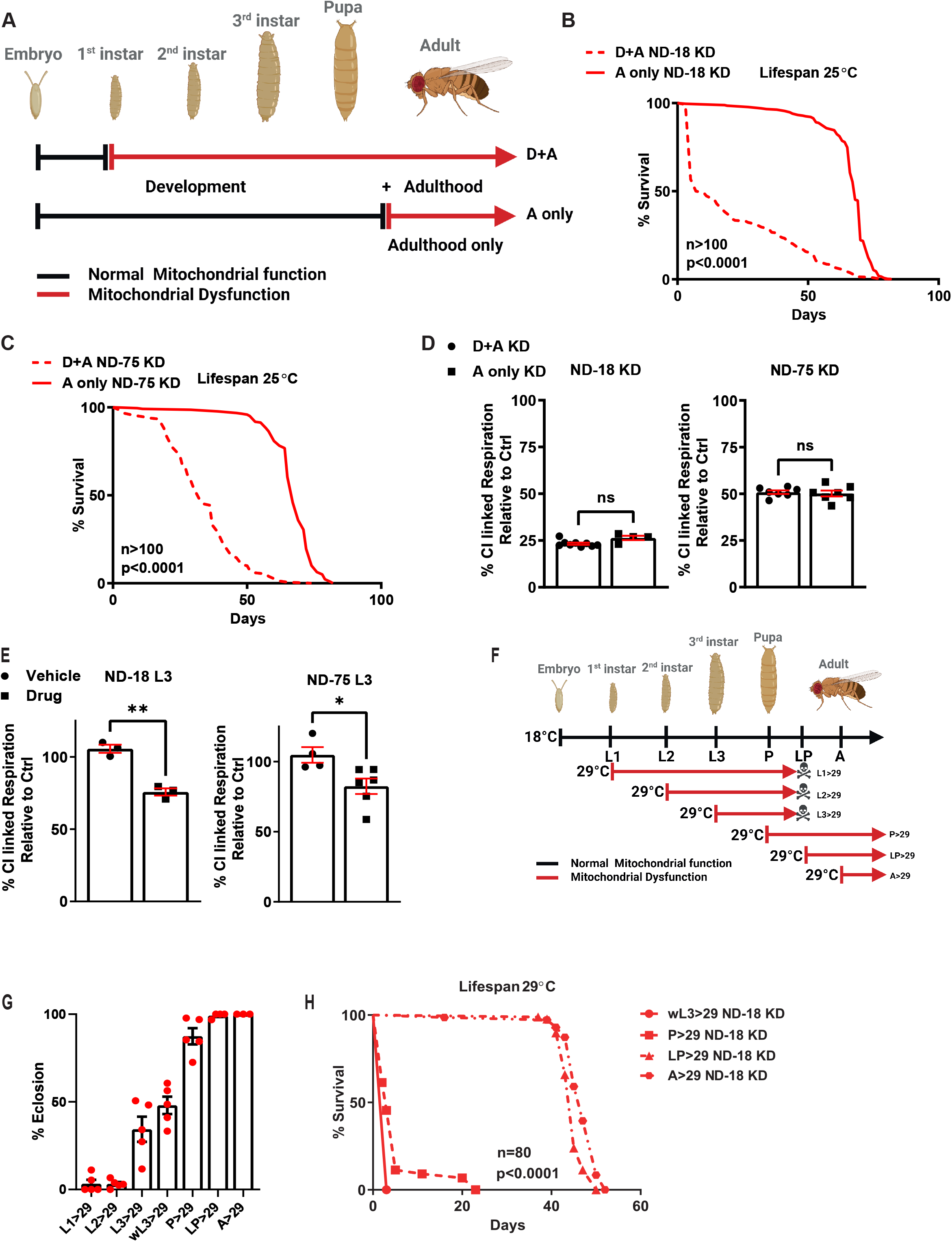
Depletion of Respiratory Complex I from development but not from adulthood shortens lifespan. (A) Schematic illustrating the periods during which the GS inducible expression system was used to reduce Complex I function, from early development into adulthood (D+A) or from Adulthood only (A only). (B) Survival of male flies where CI subunit, ND-18, has been depleted from early development (D+A) or from Adulthood (A only). (C) Survival of male flies where CI subunit, ND-75, has been depleted from early development (D+A) or from Adulthood (A only). (D) Normalised levels of CI linked respiration in ND-18 KD (left) and ND-75 KD (right) 5–7 day adult males. (E) Normalised levels of CI linked respiration in ND-18 KD (left) or ND-75 KD (right) male 3 Instar larvae in the presence (drug) or absence (vehicle) of the GS inducer, RU486. (F) Schematic depicting the developmental stages that the UAS/GAL4 expression system with the temperature sensitive form of the transcriptional repressor GAL80 (GAL80ts) expression system was used to reduce Complex I function. (G) % eclosion of flies where depletion of CI subunit, ND-18, has been induced at distinct developmental stages. (H) Survival of male flies where depletion of CI subunit, ND-18, has been induced at distinct developmental stages.

To exclude that the stark differences we observed in survival between flies where CI dysfunction had been induced from development (D+A) or from adulthood (A only) were a result of differences in CI activity, we measured levels of CI linked respiration. There was no difference in normalised CI respiratory capacity between D+A and A only KD flies for either ND-18 KD or ND-75 KD (Figure 1D, data normalised to Figure S1H). When we assayed respiration in late-stage larvae (L3) we observed, as expected, decreased CI linked respiration in ND-18 KD and ND-75 KD larvae on drug food but not those on vehicle food (Figure 1E). Notably, the level of remaining CI activity (25% ND-18 KD, 50% ND-75 KD) only correlated with lifespan in flies where CI dysfunction had been induced from development. While in flies where CI dysfunction was induced during adulthood (A only) no difference in lifespan was observed between flies with ND-18 KD, ND-75 KD, or Ctrl flies despite this large reduction in mitochondrial function, suggesting that a decrease in CI function alone is not sufficient to reduce lifespan. These data suggest that the reported loss in CI levels and activity in aged animal models and humans (Cabre et al., 2017; Navarro and Boveris, 2004; Scialo et al., 2016) may not contribute in itself to ageing or the onset of age-related diseases.

We had established that development was the window during which adaptation to mitochondrial dysfunction correlates with reduced fitness. To further narrow this down we replaced the GeneSwitch system with Gal4/Gal80 (Lee and Luo, 1999). Using ubiquitously expressed daughterless Gal4 in combination with tubulinGal80 we used two culture temperatures, 18°C (Off) and 29°C (On) to control expression of the RNAis and therefore induction of CI dysfunction more precisely (Figure 1F). Developing cultures were moved from 18 °C to 29 °C at different stages of development (Figure 1F). Eclosion data from these cultures clearly shows that lethality of CI depletion correlates with developmental timing (Figure 1G), where the earlier in development that ND-18 KD flies are shifted to 29°C the greater the level of lethality. This also correlated with survival, lifespan of flies that survived to eclosion was severely reduced proportionally and only in those flies where CI dysfunction was induced during development (Figure 1H). When the cultures were moved to 29 °C at the late pupal (LP) or eclosed adult stage no effect on lifespan was observed (Figure 1H). The lifespan of Ctrl flies subjected to the same treatments was unaffected allowing us to discount culture temperature (Figure S1I and J).

### CI depletion does not cause major developmental defects

To investigate how early vs late CI dysfunction was manifesting as reduced survival despite an equal reduction in mitochondrial function, we performed a series of physiological experiments.

We first asked if the differential effects of inducing CI dysfunction from development or from adulthood were the result of a developmental abnormality. There were no superficial phenotypic differences in adult KD or Ctrl flies treated with drug (RU486) from development or adulthood. This was important as high doses of RU486 have been shown to cause developmental effects (Andjelkovic et al., 2016). We did observe a mild developmental delay of around 24 hours and a decrease in eclosion rate of around 10% in D+A CI KD flies compared with A only and Ctrl flies (data not shown). Negative geotaxis assays commonly known as climbing assays revealed a significant reduction in climbing ability in both ND-18 and ND-75 D+A CI KD flies (data not shown). This type of climbing deficiency is often seen in models of neurodegeneration and aging (Ali et al., 2011; Madabattula et al., 2015). To rule out neurodegeneration as an underlying cause, we performed histological analysis on fly heads (Figure 2A and S2A). We found no evidence of vacuolisation, which would have been indicative of neurodegeneration, in any of the groups analysed. We also did not observe any major structural differences between ND-18 and ND-75 KD or Ctrl D+A a and A only flies. These climbing defects are also associated with muscle weakness (Ali et al., 2011). TEM analysis of indirect flight muscle showed no disorganisation of muscle fibres in D+A CI KD flies and the visible intermyofibrillar mitochondria showed no gross aberrations in mitochondrial morphology (Figure 2B and S2B). Together these data exclude developmental defects as the principal factor driving the lifespan differences seen between flies where CI dysfunction is induced during development or adulthood. Furthermore, they demonstrate that severe depletion of mitochondrial activity in adult flies is not sufficient to elicit neurodegeneration or loss of muscle architecture (Cabirol-Pol et al., 2018; Cho et al., 2012).

**Figure 2.**
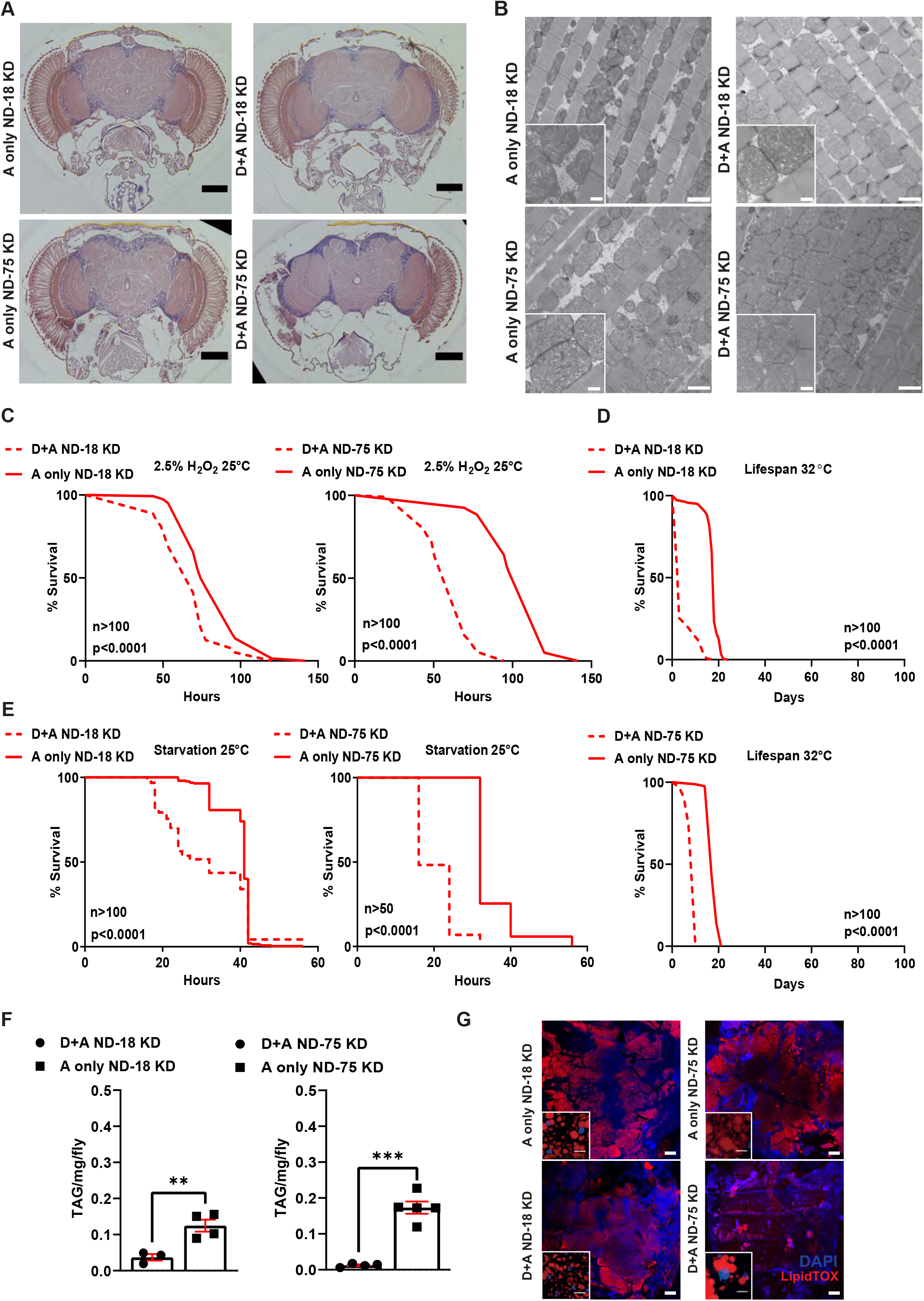
Developmental but not adulthood CI depletion increases stress sensitivity and alters energy homeostasis. (A) Representative sections of Drosophila heads from male flies where either CI subunit ND-18 (top) or ND-75 (bottom) has been depleted from early development (D+A) or from Adulthood (A only) stained with hematoxylin and eoisin, Scale bar. (B) TEM images of longitudinal sections of indirect flight muscle from males flies where either ND-18 or ND-75 has been depleted from D+A or A only, scale bar, inset scale bar (C) Survival under oxidative stress conditions of male flies where either ND-18 (left) or ND-75 (right) has been depleted from D+A or A only. (D) Survival under thermal stress conditions of males flies where either ND-18 (upper) or ND-75 (lower) has been depleted from D+A or A only. (E) Survival under starvation conditions of males flies where either ND-18 (left) or ND-75 (right) has been depleted from D+A or A only. (F) Quantification of triacylglyceride levels in males flies where either ND-18 (left) or ND-75 (right) has been depleted from D+A or A only. (G) Confocal imaging of fat bodies from male flies where either ND-18 left or ND-75 left has been depleted from D+A or A only stained with LipidTOX Red and DAPI, scale bar, inset scale bar.

### Developmental CI depletion increases stress sensitivity and alters metabolic homeostasis

Given the reduced survival of D+A CI KD flies compared with A only CI KD flies and Ctrl flies of both groups, we wanted to ask if D+A CI KD flies were short lived due to an inherent stress sensitivity. We looked at three distinct types of stress: oxidative, heat and metabolic stress. All three of these stresses should be affected by differences in mitochondrial function as they require mitochondria to adequately adapt (Graham et al., 2022; Jorgensen et al., 2021; Lewis et al., 2021) i.e., we would expect to see reduced survival in KD flies of both groups (D+A and A only) compared to Ctrl flies as they both have a 50-75% reduction in CI function. D+A and A only Ctrl flies showed no difference in stress resistance in any of the conditions suggesting that treatment with the inducer itself did not sensitise the flies to stress (Figure S2C, D and E).

Induction of oxidative stress via H_2_O_2_ treatment demonstrated that A only KD flies (ND-18 and ND-75) were as resilient to H_2_O_2_ compared to Ctrl D+A and A only flies (Figure 2C, S2C). D+A CI KD flies (ND-18 and ND-75) were significantly more sensitive, with ND-75 KD D+A flies less resistant than ND-18 D+A flies (Figure 2C, compare dashed with solid lines). Heat stress at 32°C resulted in a reduced lifespan in groups compared to the standard culture temperature of 25 °C (Figure 2D, S2D). However only flies which had CI dysfunction from development (D+A KD flies ND-18 and ND-75) were short-lived in comparison to their A only counterparts (Figure 2D, compare dashed with solid lines). Finally, we subjected all groups to starvation at 25°C. As with oxidative and heat stress, starvation resistance in A only KD flies was in accordance with what we observed for Ctrl D+A and A only flies, while D+A ND-18 and ND-75 KD flies were significantly more sensitive (Figure 2E, S2E, compare dashed with solid lines).

Whole fly histological analysis of D+A vs A only KD flies indicated a reduction in adipose tissue in adult flies, known as the fat body (data not shown). The fat body is the major lipid storage tissue, second arm of the cellular immune response in *Drosophila* and analogous to the mammalian adipose tissue and liver (Baker and Thummel, 2007). To ascertain whether the reduction of visible adipose tissue was affecting lipid storage we measured levels of whole fly triglycerides (TAG) in all groups (Figure 2F and S2F). D+A and A only Ctrl flies had similar levels of TAG, while both ND-18 KD and ND-75 KD D+A flies had significantly less TAG than the corresponding A only flies, confirming a specific decrease in flies where CI has been depleted during development. TAG levels in D+A ND-75 KD flies were almost undetectable, despite higher levels of CI function than in D+A ND-18KD flies. Dissection and staining of adult abdominal fat bodies with a neutral lipid stain, LipidTOX, further corroborated the TAG data (Figure 2G and S2G). We observed decreased levels of stored lipids in D+A CI KD flies compared with A only CI KD flies (Figure 2G). ND-75 KD D+A flies had almost no discernible adult fat body, but remnants of larval fat body attached to trachea, which would normally be fully histolysed within days of adult eclosion implying a delay in the metabolic transition to adulthood. In ND-18 KD flies we saw differences in the intensity of LipidTOX staining as well as side of lipid droplets within the fat body. This reduction in stored lipids specifically in D+A CI KD flies could underly reduced starvation resistance in comparison to A only CI KD flies (Figure 2E).

Given the levels of CI dysfunction we were inducing, it was important to understand the effect on the NAD+:NADH ratio. The NAD+:NADH ratio has been implicated in the pathology of several diseases with mitochondrial dysfunction (Liu et al., 2021; Santidrian et al., 2013; Titov et al., 2016). We detected no effect of drug treatment on levels of NAD+, NADH or the NAD+:NADH ratio in Ctrl D+A or A only flies (Figure S2H). D+A ND-75 KD flies had a slight but significant decrease in NAD+ and NADH in comparison to A only ND-75 KD flies but this was not mirrored in ND-18 KD flies (Figure S2H and I) indicating that the NAD+:NADH ratio is not contributing to the differential longevity of these flies.

This data demonstrates that CI dysfunction alone is not sufficient to reduce lifespan and increase stress sensitivity as flies where CI dysfunction is induced from adulthood are indistinguishable from Ctrl flies. We hypothesized that the induction of CI dysfunction during development, where adaptive plasticity is at its highest, causes maladaptation which results in reduced survival and stress resistance.

### Developmental but not adulthood CI depletion results in specific maladaptive response

We had established that developmental mitochondrial dysfunction resulted in decreased lifespan and increased stress sensitivity (Figure 1 and 2), while inducing the same level of CI depletion in adult flies did not elicit any of these phenotypes. We reasoned that the response to CI dysfunction during development must underlie differences in the physiological effects of CI dysfunction. We hypothesized that the inherent plasticity of development, reduced or absent in adulthood, could allow for maladaptive responses to CI dysfunction.

To test whether CI dysfunction in development vs adulthood was provoking a differential adaptive response, we first assessed the transcriptional response using RNAseq analysis of whole adult flies from all groups (Figure S3A). Principal component analysis implied the expression profiles of D+A vs A only KD flies were distinct (Figure S3B and C), while for Ctrl flies there was significantly more overlap (Figure S3D). Comparing gene expression in Ctrl D+A vs A only flies identified a small set of significantly altered genes (Figure S3E). Treatment with various concentrations of RU486 has been reported to alter gene expression in several fly models. To remove any effect on gene expression that might be the result of the RU486 treatment regime used to induce CI dysfunction, we used an inhouse bioinformatic pipeline to filter out those genes differentially regulated in Ctrl D+A vs A only from our KD expression data (Figure S3A and E). CI dysfunction induced from development vs adulthood (D+A vs A) with KD of either ND-18 or ND-75 resulted in the significant differential expression of many genes (Figure 3A and B). ND-75 KD flies showed a greater response than ND-18 KD flies. Although, we saw upregulation of some genes, most genes were overwhelmingly downregulated. Importantly, comparative analysis of gene expression changes between ND-18 KD D+A vs A only and ND-75 D+A vs A only, demonstrated a high level of concordance (Figure 3A, B and C, significant genes in pink, significant genes in both comparisons in red). Establishing that the response to CI dysfunction in both models overlaps highly.

**Figure 3.**
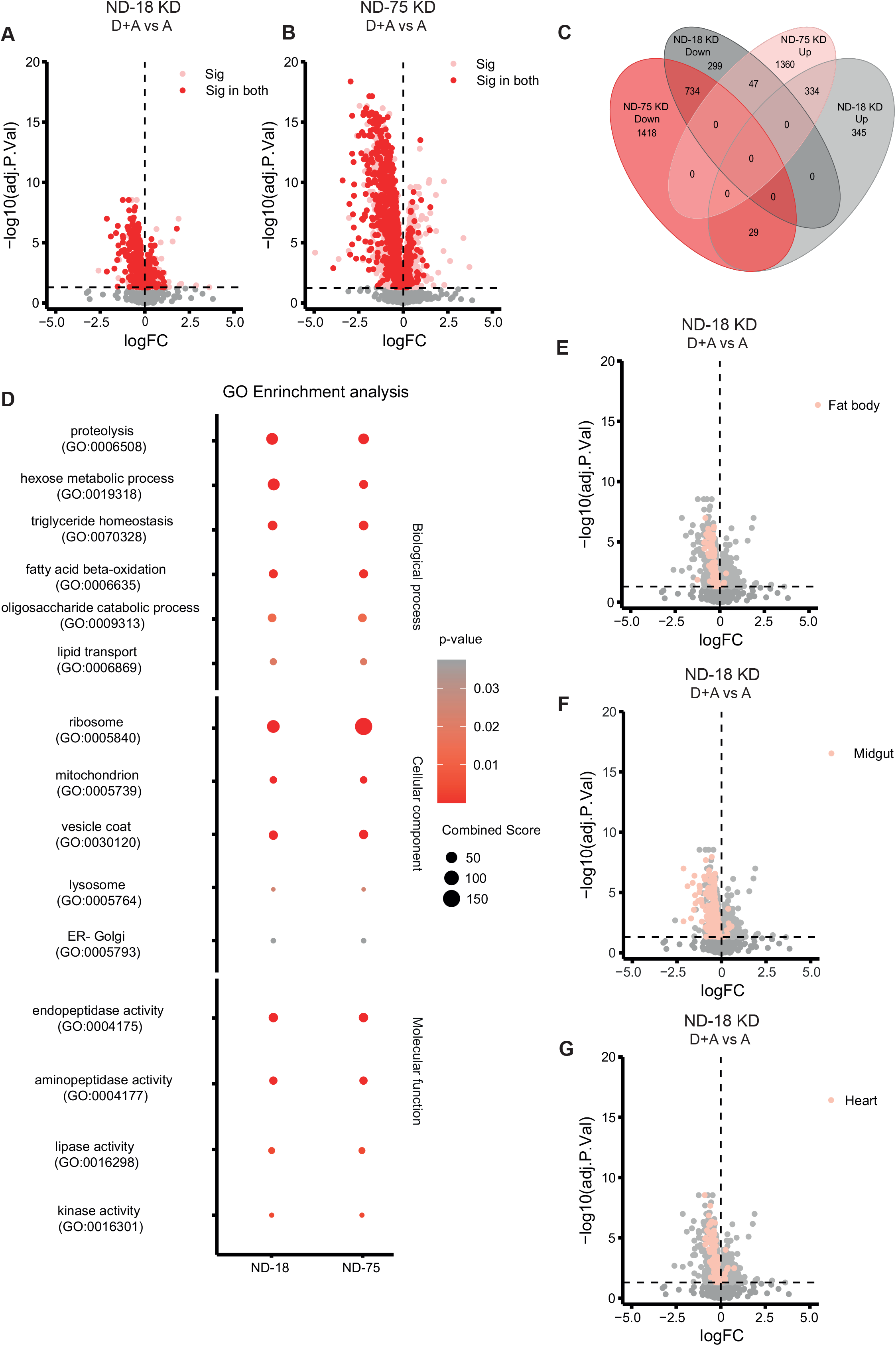
Developmental CI depletion results in a specific transcriptomic signature. (A+B) Volcano plot of genes differentially expressed between flies where ND-18 (A) or ND-75 (B) was depleted from early development (D+A) vs from Adulthood only. Each dot represents a single gene. In pink are those genes that are significantly altered, in red are those which are significantly altered in both ND-18 and ND-75 D+A KD flies. In grey are genes which were not significantly altered. (C) Venn diagram illustrating the relationship between the differential expression seen in ND-18 D+A vs A and ND-75 D+A vs A. (D) GO enrichment analysis (FlyEnricher) performed on those genes significantly altered in both ND-18 D+A vs A and ND-75 D+A vs A reveals significant enrichment in the following classifications. (E, F & G) Volcano plot of genes significantly differentially expressed between ND-18 D+A and ND-18 A (grey), highlighted in pink are those genes whose expression is known to be highly enriched in the fat body (E), midgut (F) and heart (G).

Gene Ontology (GO) Enrichment Analysis of the genes significantly differentially expressed in both ND-18 and ND-75 D+A KD flies revealed a transcriptional signature characterized by dysregulated lipid homeostasis, transport and metabolism (GO:0070328, GO:0006869, GO:0006635, GO:0016298), reduced protein and sugar metabolism (GO:0006508, GO:0019318, GO:0009313, GO:0004175, GO:0004177, GO:0005793, GO:0005764, GO:0030120) and translation (GO:0005840) (Figure 3D). We used clustergrams to look at the expression of all genes associated with fatty acid beta oxidation, triglyceride homeostasis, mitochondria, and protein synthesis (Figure S3J and K). We found that most genes associated with each of GO terms were differentially regulated in our D+A CI KD flies. This common signature in D+A CI KD flies reinforced a suppression of both catabolic and anabolic metabolism in response to developmental CI dysfunction. KEGG pathway analysis of further confirmed this (Figure S3F), with sugar, lipid and amino acid metabolism most significantly affected. This data is also congruent with our findings of reduced lifespan, increased stress sensitivity and dysregulation of lipid homeostasis.

We chose to deplete CI ubiquitously as often is the case in patients, but we know that mitochondria are enriched in tissues with higher energy demands, such as the skeletal muscle, heart, and brain. This is very clear when we look at the clinical phenotypes associated with mitochondrial dysfunction (Gorman et al., 2016). Already in our own models presented here, we have observed alterations in the fat body in flies where CI dysfunction is induced from development (Figure 2G, S2G). We used tissue RNAseq enrichment data from Flyatlas2 (Leader et al., 2018) to see whether a specific tissue was contributing to the shared transcriptional signature of D+A CI KD flies. We identified major contributions from three tissues, the fat body (Figure 3E and S3G), midgut (Figure 3F and S3H) and heart (Figure 3G and S3I). The fat body and midgut are essential to management of organismal energy balance. Several studies have demonstrated the role of mitochondrial metabolism in the proper differentiation and function of the intestine (Deng et al., 2018; Liu et al., 2016; Wisidagama and Thummel, 2019; Zhang et al., 2022), as well as its requirement in the fat body for long range growth control (Banerjee et al., 2013; Song et al., 2017). In humans the myocardium is the richest source of mitochondria (Page and McCallister, 1973) and in flies proper mitochondrial dynamics are essential for cardiac function (Dorn et al., 2011). Our data imply that although there is a ubiquitous decrease in CI function, the maladaptive response in D+A KD flies is most acute in the fat body, midgut, and heart.

Mitochondrial dysfunction is well known to result in metabolic alterations, in fact many of the most effective interventions published that prevent or attenuate these alterations target metabolic dysfunction (Russell et al., 2020). To investigate if the distinct transcriptomic signature of D+A vs A only CI-KD flies extended to the metabolome, we performed whole fly metabolomics. Treatment with RU486 during development resulted in an expected minor, if significant metabolomic response in Ctrl flies (Ctrl D+A vs A) (Figure S4A, B and C). As with our transcriptomic analysis, to remove any effects specific to RU486 treatment any metabolites found to be significantly altered in Ctrl flies were removed from our analysis of CI-KD flies (ND-18 KD and ND-75 KD flies). Comparison of the metabolomes of D+A vs A only ND-18 KD flies (Figure 4A and S4D) revealed a significant number of altered metabolites. This was notably escalated in D+A vs A only ND-75 KD flies where we detected a larger number of significantly altered metabolites (Figure 4B and S4E), mirroring what we have seen regarding lipid storage and the transcriptome. Using the MS Peaks to Pathways module (Pang et al., 2021) of MetaboAnalyst we performed functional enrichment analysis. We did this separately for ND-18 and ND-75 KD flies. Despite the disparity we saw in the number of altered metabolites between ND-18 and ND-75 KD flies there was large level of convergence amid the metabolic pathways significantly altered in D+A vs A only KD flies. Most notably, like in our transcriptomic analysis, we found sugar, lipid, and amino acid metabolism to be most significantly affected (Figure 4C) in short lived D+A vs long lived A only CI KD flies, further supporting a role for metabolic plasticity in determining the effects of mitochondrial dysfunction and our hypothesis that this plasticity facilitates maladaptive responses that are detrimental to organismal fitness.

**Figure 4.**
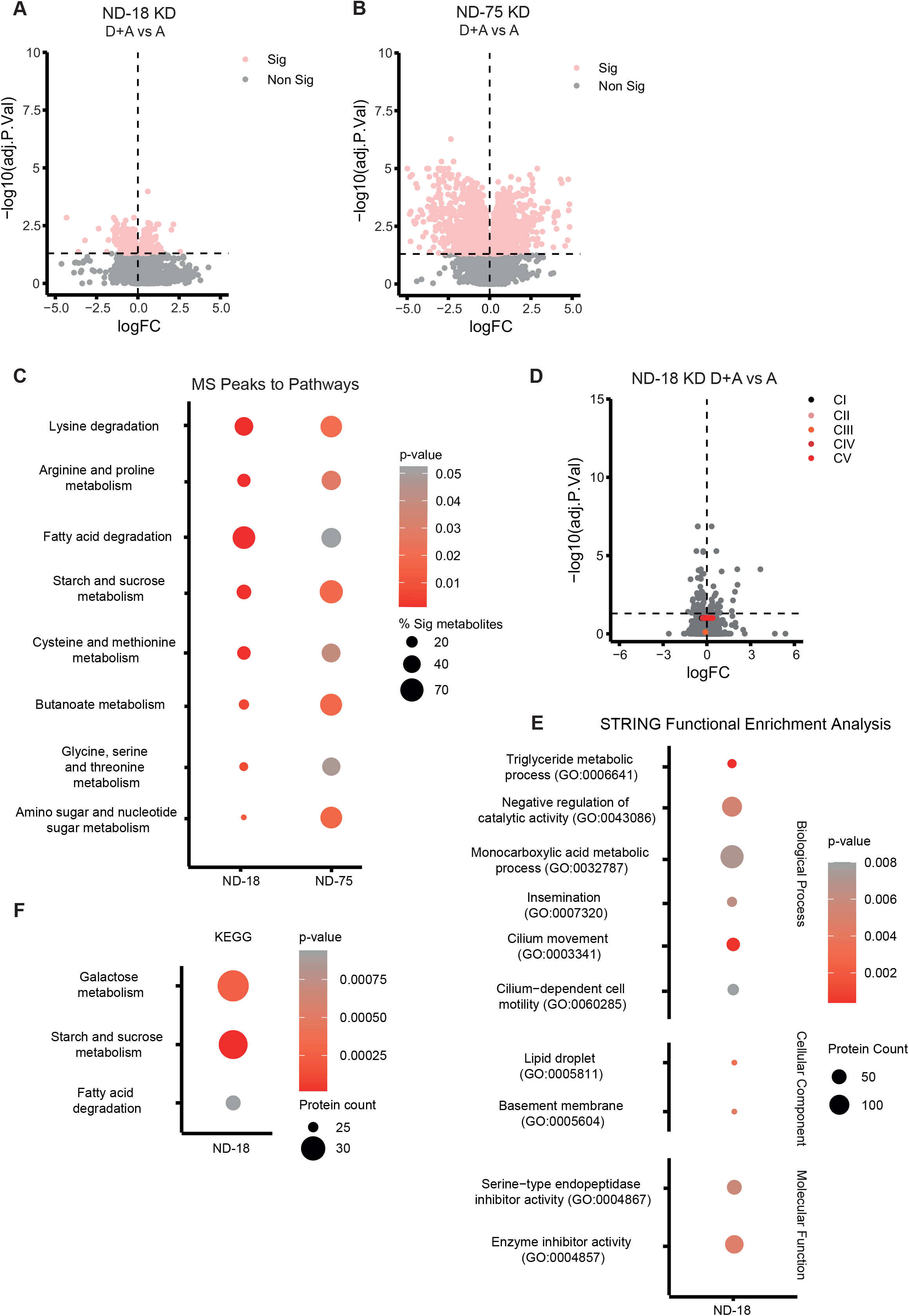
Developmental CI depletion results in a specific metabolomic signature. (A+B) Volcano plot of metabolites with significantly different abundance between flies where ND-18 (A) or ND-75 (B) was depleted from early development (D+A) vs from Adulthood only (A only). (C) Functional enrichment analysis using MetaboAnalyst MS Peaks to Pathways (KEGG) of the differential metabolic profiles of ND-18 and ND-75 KD D+A vs A only, identified enrichment of the following categories. (D) Volcano plot of proteins with significantly different abundance between flies where ND-18 has been depleted from development (D+A) vs Adulthood only (A only), with subunits of the five ETC complexes highlighted (E) Dot plot showing the results of STRING Functional Enrichment Analysis of ND-18 KD D+A vs A only flies (F) Dot plot showing the most significant pathways (KEGG) altered at the protein level in ND-18 KD D+A flies.

Finally, we wanted to ask if we could detect any changes in the proteome of our short-lived CI depleted flies (Figure S4F and G). Proteomic analysis of whole fly homogenates confirmed depletion of CI subunits in D+A and A only ND-18 KD flies (Figure S4A, H and I), while subunits of CII, III, IV and V were unaffected. Further, we detected no differences in levels of ETC subunits between D+A flies and A only ND-18 KD flies (Figure 4D), confirming again that the differential effects on lifespan, gene expression and metabolism were not a result in differing levels of ETC function. Using STRING we performed functional enrichment analysis of proteins found to have significantly different abundance in D+A vs A only ND-18 KD (Figure 4E, F and S4J). Like our transcriptomic and metabolomic analysis of flies where CI dysfunction has been depleted from development, this functional enrichment analysis highlighted a signature of dysregulated anabolic and catabolic lipid, protein, and sugar metabolism.

## Conclusion

We have developed unique *in vivo* models in which we have temporally manipulated the function of CI. Our molecular and physiological characterisation of short- and long-lived models of CI dysfunction establish development as the phase at which levels of mitochondrial dysfunction can determine lifespan and organismal fitness. Bringing light to an area of great contradiction in the field regarding the role of CI in longevity and implies that treatments targeting mitochondrial dysfunction should be targeted during this critical window. Our data identify and detail the unique maladaptive responses to developmental CI dysfunction that are absent in long lived flies with equal levels of CI dysfunction, which could be targeted therapeutically. We have identified the key tissues making the largest contributions to the response to mitochondrial dysfunction and lifespan. Our data demonstrating that almost 80% of CI function is dispensable in adulthood also invite us to rethink the role of CI function in the aging process. To realise the potential of this work we must now focus on understanding how and which of these adaptations we are able to target to prevent and reverse the effects of mitochondrial dysfunction in disease and aging.

## Acknowledgments

This research was supported by a Sir Henry Wellcome Postdoctoral Fellowship to R.S (204715/Z/16/Z), a Wellcome Senior Research Fellowship (212241/A/18/Z) and BBSRC grants (BB/R008167/1 & BB/W006774/1) to A.S and fellowships from the Uehara Memorial Foundation and International Medical Research Foundation to T.K.. The work of the lab of L.M.M. is funded by the UK Medical Research Council intramural project MC_UU_00025/3 (no. RG94521). We thank the Vienna and Bloomington Drosophila Stock Centres for fly stocks. We thank Colin Nixon (Beatson CRUK histology service) and Julia Cordero (UofGlasgow) for advice on whole fly immunohistochemistry. We thank the Newcastle University Electron Microscopy Service for their help with the transmission electron microscopy. We thank D. McGuinness and J. Galbraith (UofGlasgow, Polyomics) for processing RNAseq samples. We would like to thank Dr Catarina Franco and Dr Bini Ramachandran for help with developing the proteomic sample processing pipeline, sample preparation and data processing, as well as providing helpful discussions on sample analysis and experimental design. We thank all past and present members of the Sanz lab for useful discussion and technical support.

## Author Contributions

R.S conceived and co-supervised the project, designed, carried out most experiments and analysed and interpreted the data. F.R processed and analysed all RNAseq, metabolomic and proteomic data. A.H.U and O.M performed metabolomics analysis. K.M collected and prepared samples for proteomics analysis. B.C.A performed H2O2 stress analysis. Y.Y and L.M.M performed proteomics analysis. T.K and V.I.K performed NAD+/NADH analysis. R.S wrote the first draft of the paper with contributions from the rest of the authors. A.S co-supervised the project, designed experiments, analysed, and interpreted data.

## Ethical Statement/Conflict of Interest

V.I.K is a Scientific Advisor for Longaevus Technologies. Other authors declare no competing interests.

---

**Figure S1.**
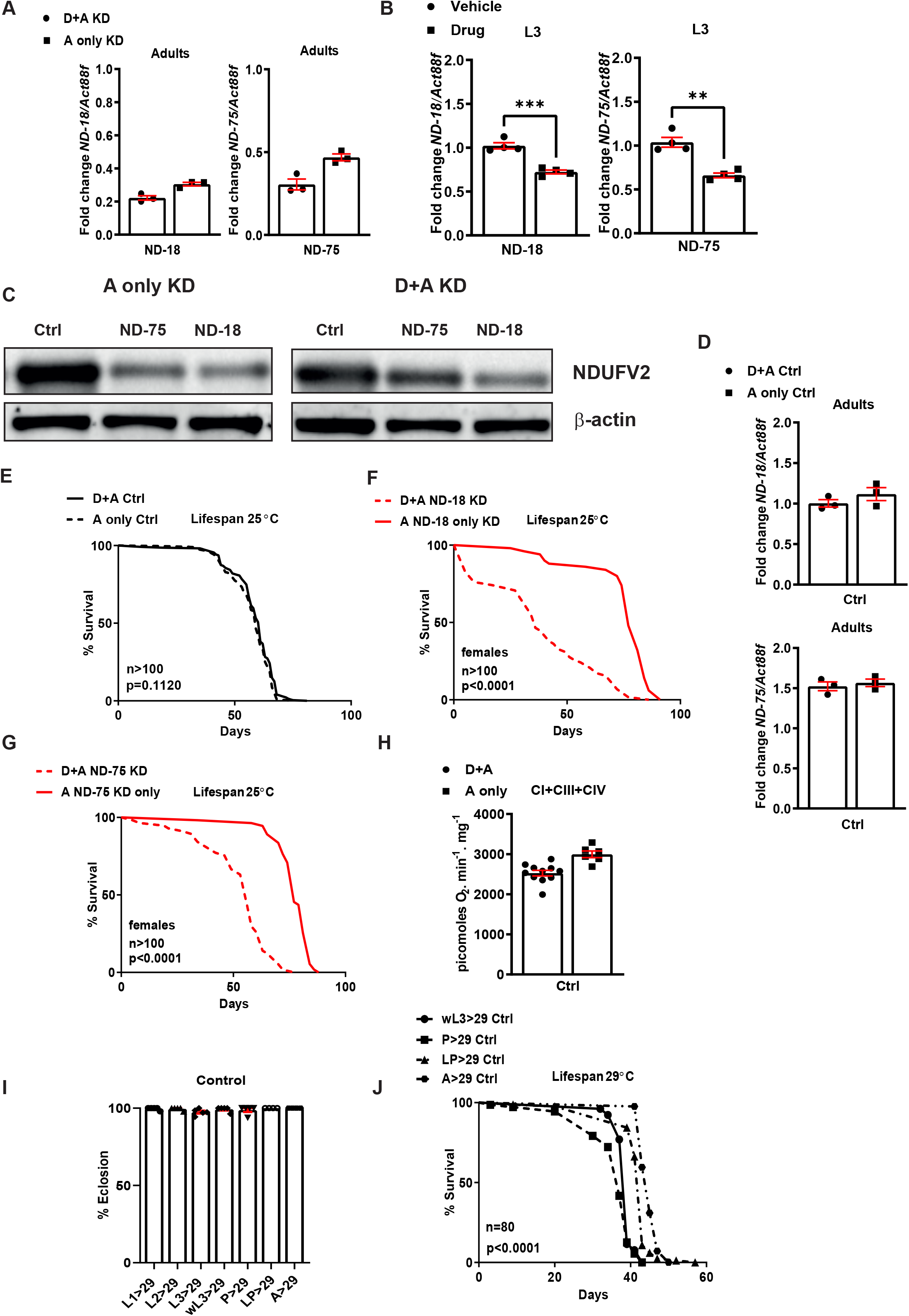
(A) Quantification of KD in ND-18 KD (left) and ND-75 KD (right) 5–7 day adult males. (B) Quantification of KD in ND-18 KD (left) and ND-75 KD (right) 3^rd^ instar larvae in the presence (drug) or absence (vehicle) of the GS inducer, RU486. (C) Western blot analysis of CI core subunit, NDUFV2 (ND-24) in control males and males where ND-18 or ND-75 has been depleted from early development (D+A KD) or from adulthood only (A only KD), beta-actin was used as a loading control. (D) Quantification of ND-18 (upper) and ND-75 (lower) mRNA levels in 5–7 day adult males exposed to the inducer from D+A or A only (E) Survival of male control flies exposed to the inducer, RU486 from early development (D+A) or from Adulthood (A only). (F) Survival of female flies where CI subunit, ND-18, has been depleted from early development (D+A) or from Adulthood (A only). (G) Survival of female flies where CI subunit, ND-75, has been depleted from early development (D+A) or from Adulthood (A only). (H) Levels of CI linked respiration in control 5–7 day adult males exposed to the inducer from D+A or A only. (I) % eclosion of control males flies which have been moved from 18°C to 29°C at distinct developmental stages. (J) Survival of control male flies which have been moved from 18°C to 29°C at distinct developmental stages.

**Figure S2.**
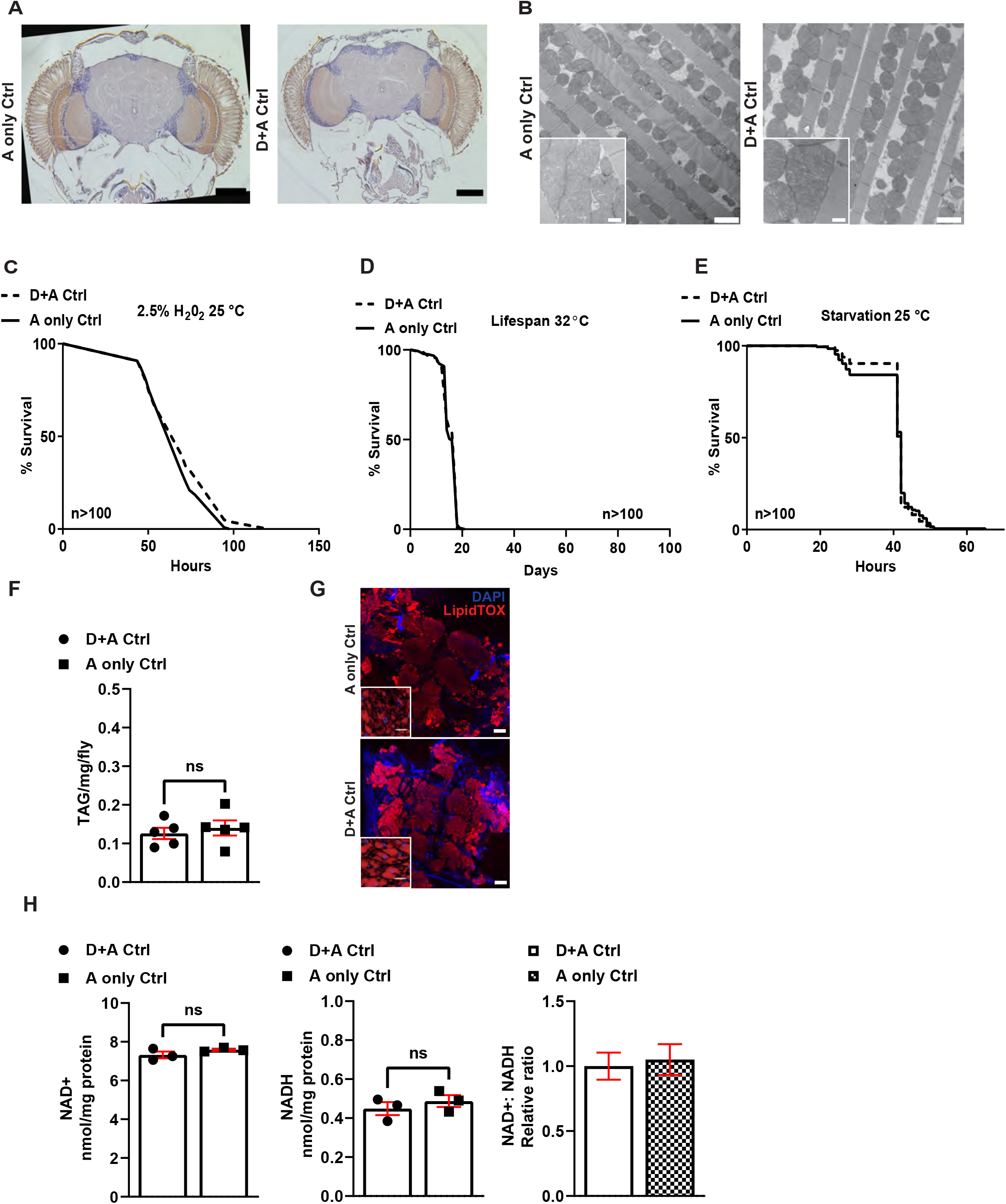
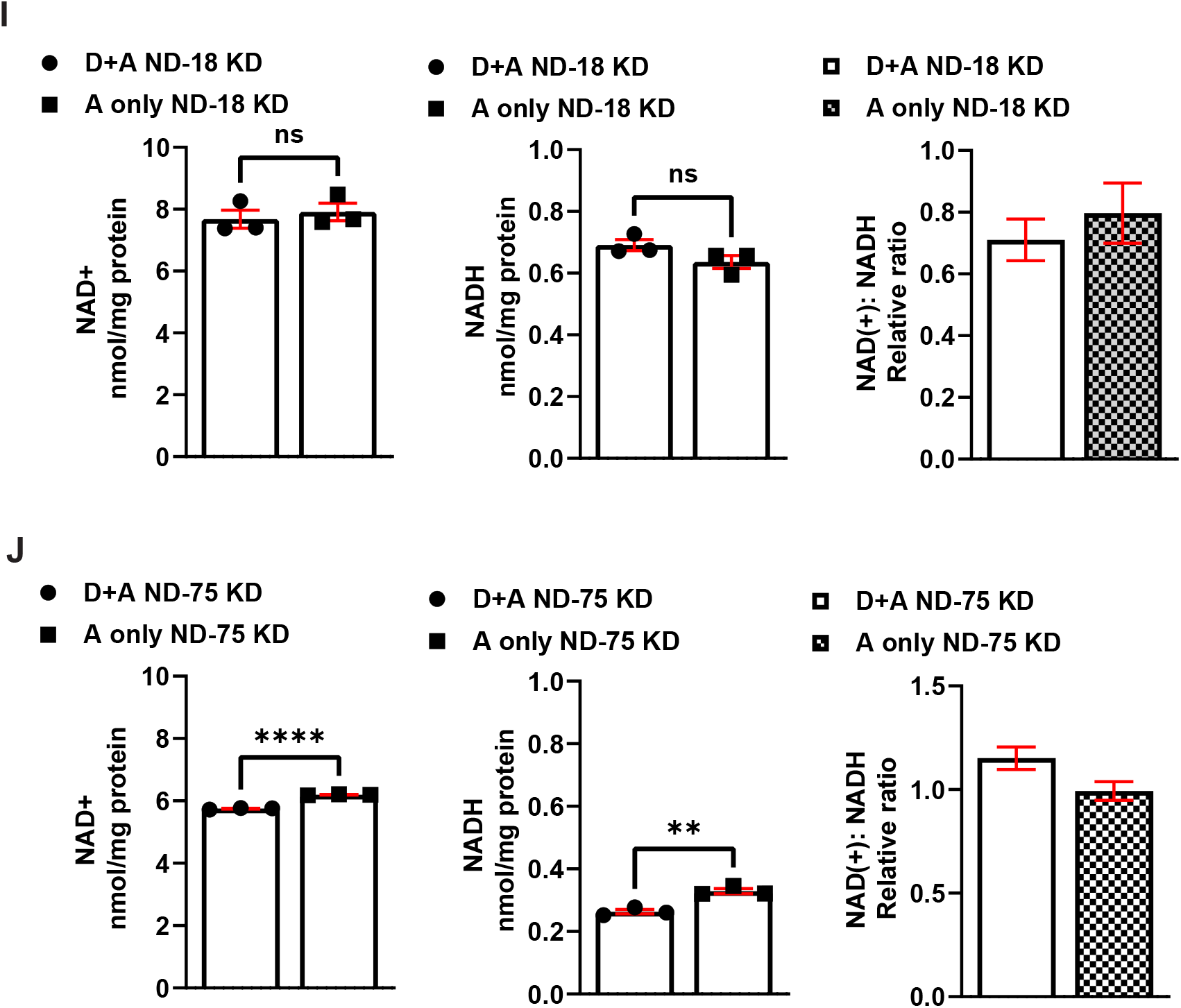
(A) Representative sections of Drosophila heads from control male flies exposed to the inducer, RU486 from early development (D+A) or from Adulthood (A only) stained with hematoxylin and eoisin, Scale bar, (B) TEM images of longitudinal sections of indirect flight muscle from control males flies control male flies exposed to the inducer, RU486 from early development (D+A) or from Adulthood (A only), scale bar, inset scale bar. (C) Survival under oxidative stress, (D) thermal stress and (E) starvation conditions of control males flies exposed to the inducer, RU486 from early development (D+A) or from Adulthood (A only). (F) Quantification of triacylglyceride levels in control males flies exposed to the inducer, RU486 from early development (D+A) or from Adulthood (A only). (G) Confocal imaging of fat bodies from male flies exposed to the inducer, RU486 from early development (D+A) or from Adulthood (A only) stained with LipidTOX Red and DAPI, scale bar, inset scale bar. (H, I and J) Levels of NAD+, NADH and the relative ratio of NAD(+):NADH in control (H), ND-18 KD (I) and ND-75 KD male flies (J).

**Figure S3.**
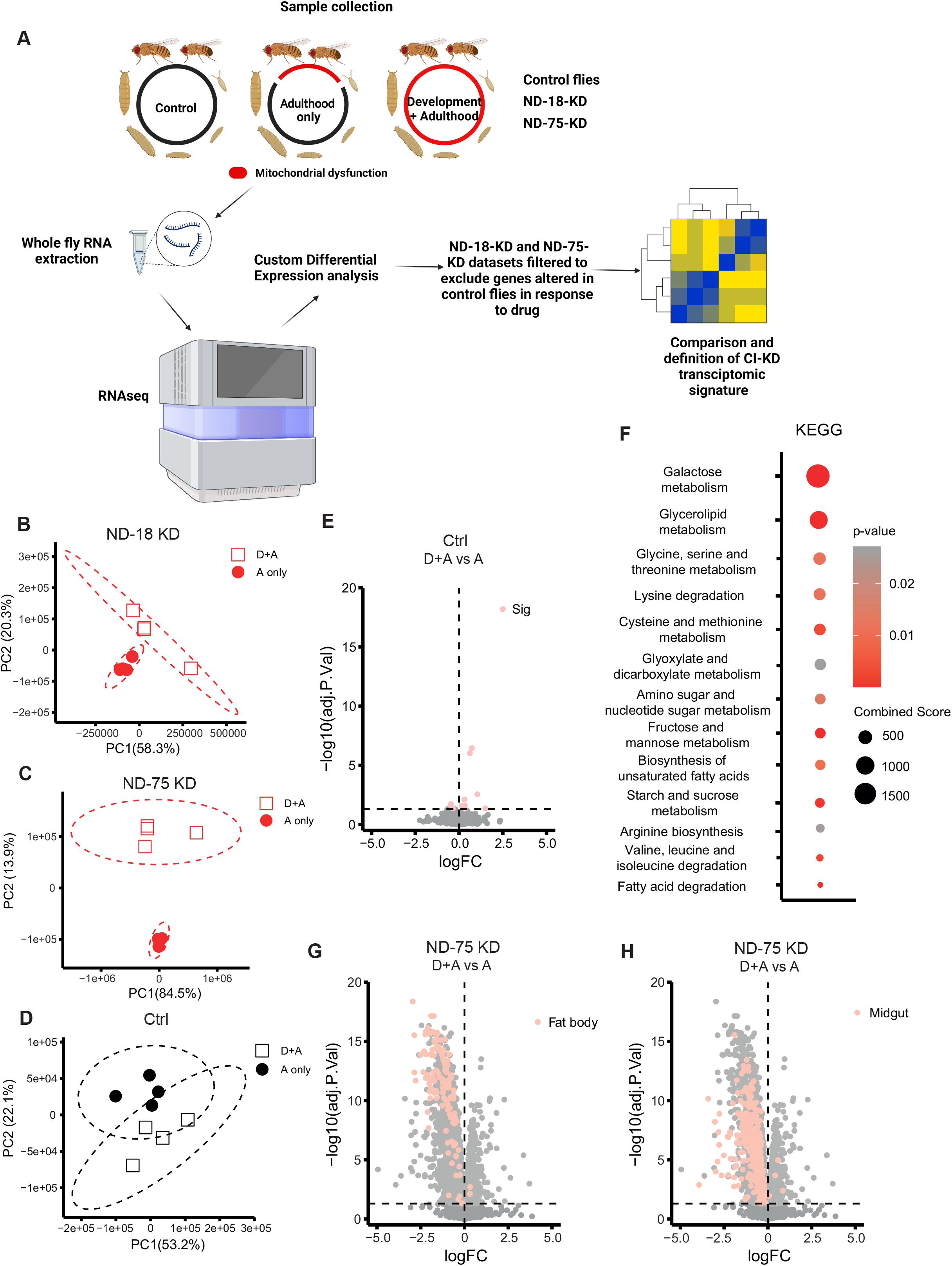
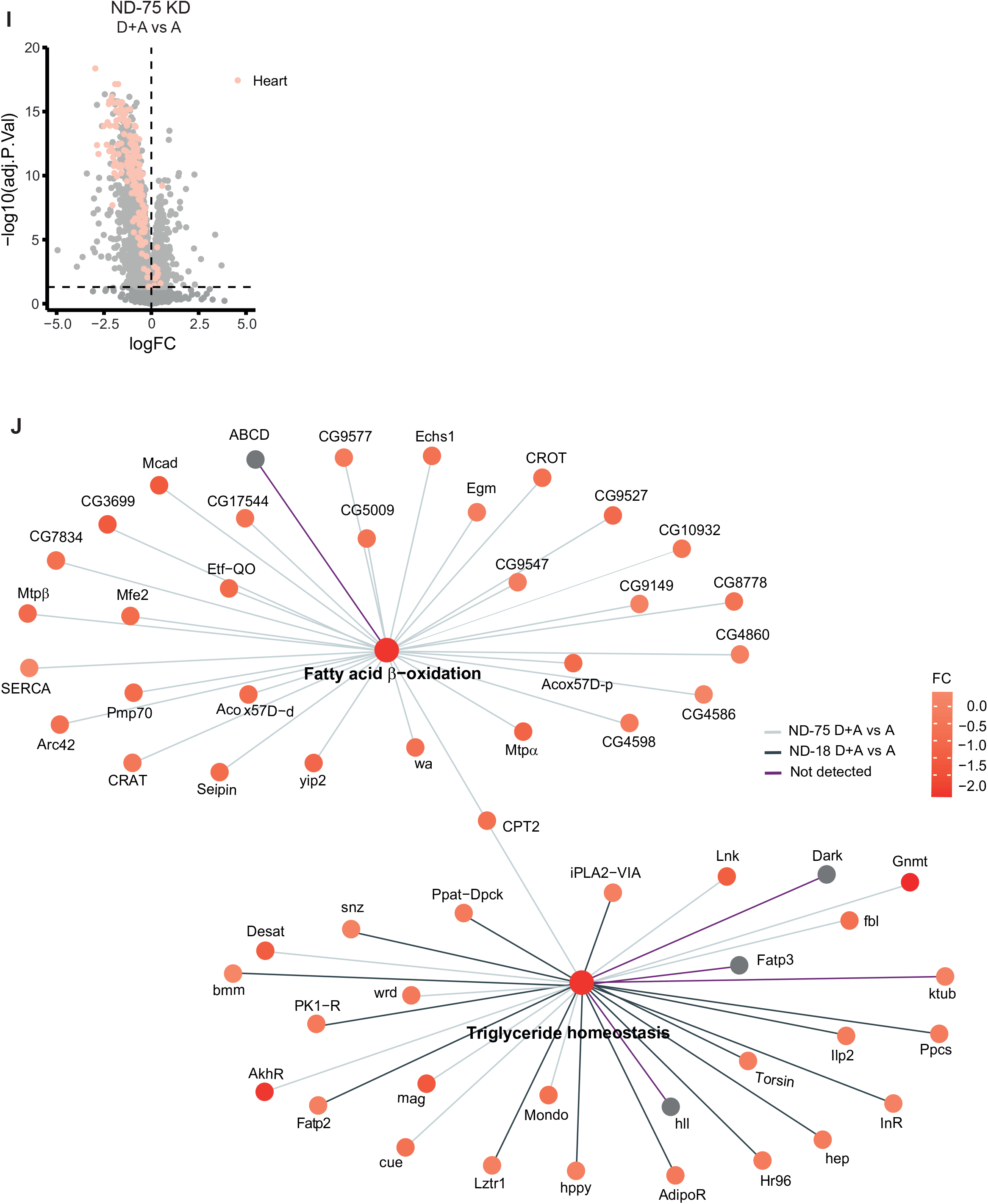
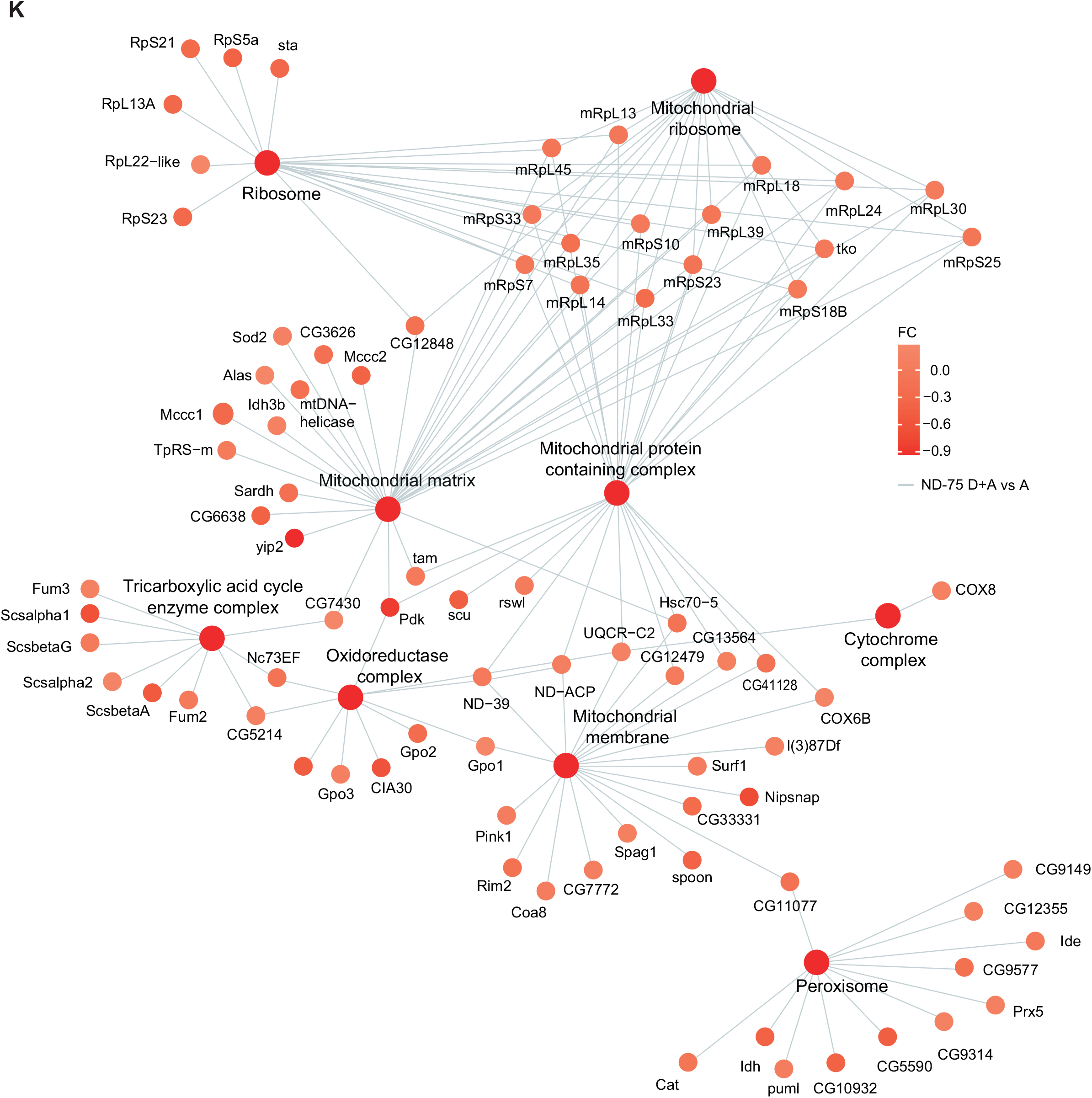
(A) Schematic representation of data analysis pipeline used to collect, process, and analyse transcriptomic data. (B, C & D) PCA analysis of ND-18 KD (B) and ND-75 KD (C) and Ctrl (D) D+A vs A conditions. (E) Volcano plot of genes significantly differentially expressed between Ctrl D+A and Ctrl A. (F, G & H) Volcano plot of genes significantly differentially expressed between ND-75 D+A and ND-75 A (grey), highlighted in pink are those genes with highly enriched expression in the fat body (F), midgut (G) and heart (H). (I & J) Clustergrams of the indicated GO terms expressing the differential expression (FC) of genes associated with this term. All FCs are from ND-75 KD D+A vs A only comparison unless otherwise indicated by a dark grey line (ND-18 KD D+A vs A only) or a purple line (not detected in our study).

**Figure S4.**
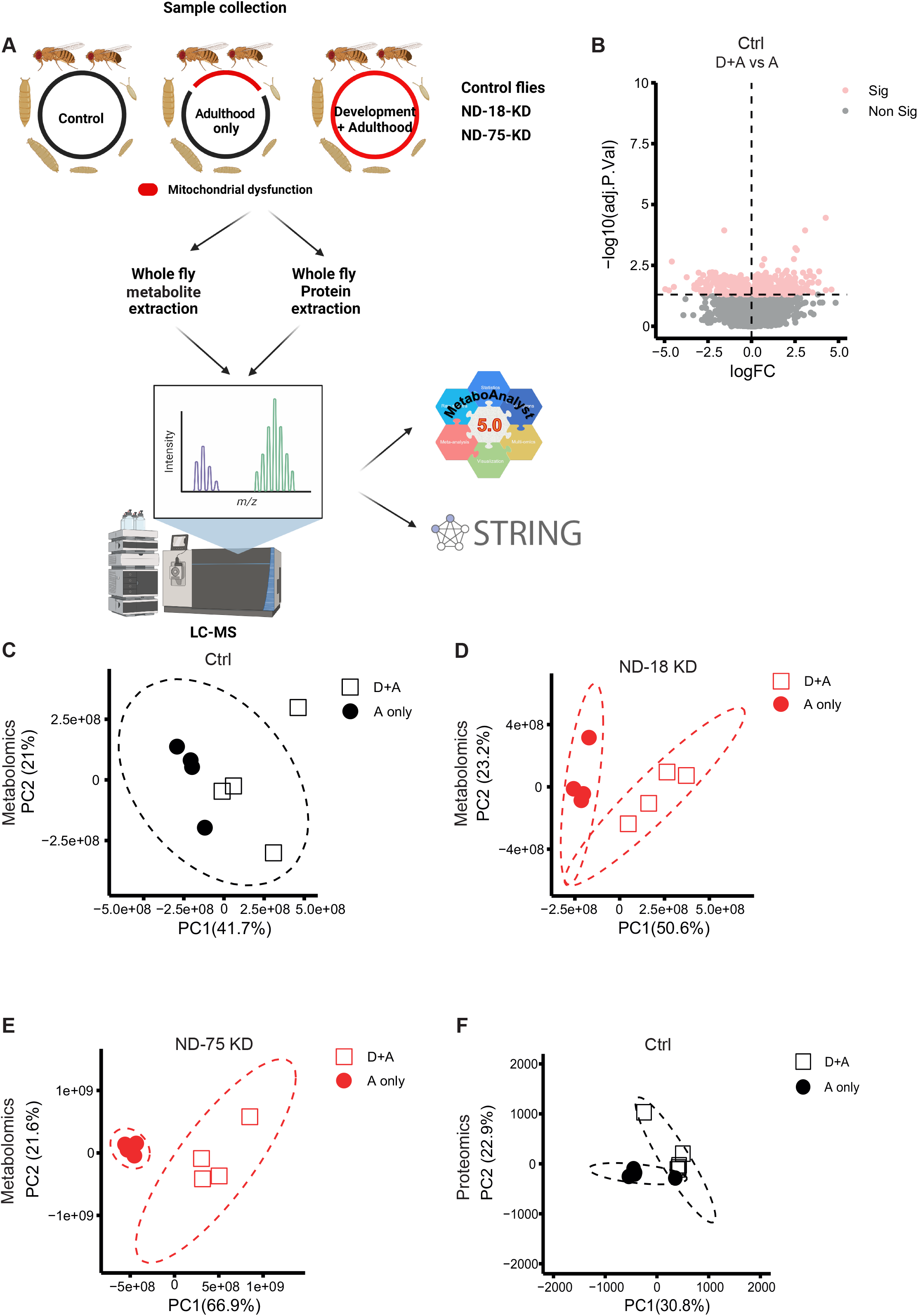
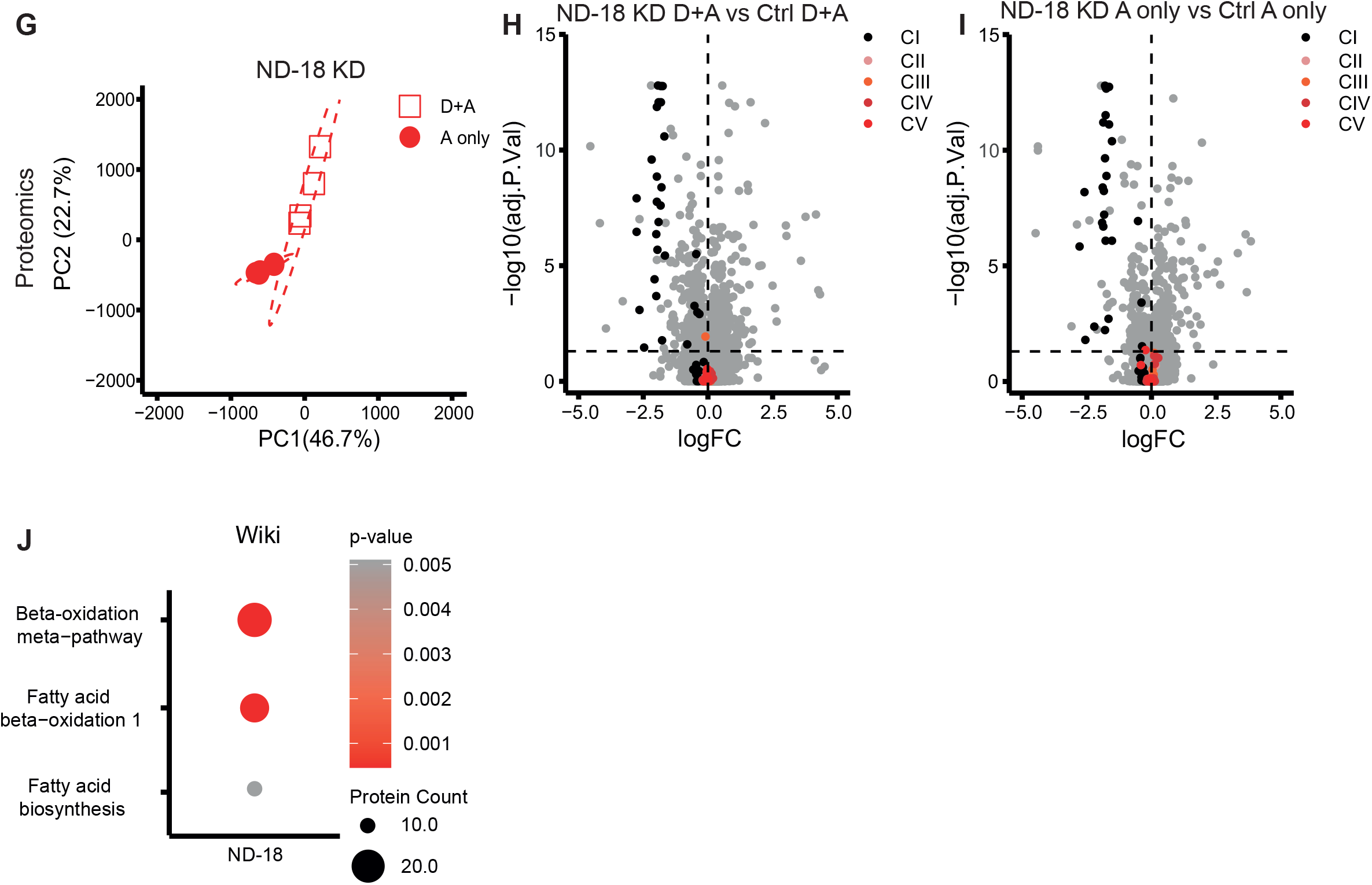
(A) Schematic representation of data analysis pipeline used to collect, process, and analyse metabolomic and proteomic data. (B) Volcano plot of metabolites with significantly different abundance between Ctrl D+A and Ctrl A. (C, D & E) PCA analysis of metabolomics data from Ctrl (C) and ND-18 KD (D) and ND-75 KD (E) D+A vs A conditions. (F & G) PCA analysis of proteomics data from Ctrl (F) and ND-18 KD (G) D+A vs A conditions. (H) Volcano plot of proteins with significantly different abundance between flies where ND-18 has been depleted from development (D+A) vs Ctrl D+A flies, with subunits of the five ETC complexes highlighted (I) Volcano plot of proteins with significantly different abundance between flies where ND-18 has been depleted from Adulthood (A only) vs Ctrl A only flies, with subunits of the five ETC complexes highlighted (J) Dot plot showing the most significant pathways (Wiki) altered at the protein level in ND-18 KD D+A flies

