## Supplementary materials for "Developmental mitochondrial Complex I activity determines lifespan"

**Materials and Methods**

**Genotypes**

**Supplementary Table 1. Genotype and source information for fly lines used in this study**.

| Genotypes | Source |
| --- | --- |
| *w^1118^;; tubulinGeneSwitch* (*tubGS*) | Gifted by Dr Scott Pletcher |
| *w^1118^;; daughterlessGal4*  (*daGal4*) | BDSC |
| *w^1118^;tubulinGal80^ts^*  (*tubGal80^ts^*) | BDSC |
| *w^1118^; UAS-ND-18-IR*  (*ND-18-KD*) | VDRC, CG12203, 101489/KK |
| *w^1118^; UAS-ND-75-IR*  (*ND-75-KD*) | VDRC, CG2286, 100733/KK |

Lines were backcrossed into *w^1118^* background for 11 generations

**Supplementary Table 2. Primer Sequences**

| Primer | Sequence |
| --- | --- |
| *Act88f* F | agggtgtgatggtgggtatg |
| *Act88f* R | cttctccatgtcgtcccagt |
| *ND-18* F | aagatcaccgtgccgactg |
| *ND-18* R | gacaatgggtcgccgctg |
| *ND-75* F | tgggagatccgtaaggtgag |
| *ND-75* R | ctcgttgacgtcctcgttct |

**in-house RNA sequencing pipeline**

Raw reads are pre-processed through TrimGalore V.0.6.5, which combines FastQC to determine quality and Cutadapt (Martin, 2011) to trim reads if necessary. Both the adaptive quality and adapter trimming were employed within the script. Mapping of the trimmed reads was conducted in HISAT2 V.2.2.1 (Kim et al., 2019). All reads were aligned to the pre-built Hisat2 D. Melanogaster BDGP6 index (obtained through HISAT2’s website – last accessed 14/9/22). The resultant Sam files were sorted and converted to Bam files using Samtools V.1.10 (Danecek et al., 2021), then counts were then established using StringTie V.2.1.1 (Pertea et al., 2016). Each sample was annotated via StringTie using ‘dmel-all-r6.41.gtf’ (available from Flybase) as a reference. All stages, as well as several intermediate stages with file type conversions, were connected to custom Python3 scripts which are available on request.

**Supplementary Table 3. R. Packages**

| Package Name | Version | Link |
| --- | --- | --- |
| dplyr | 1.0.10 | <https://dplyr.tidyverse.org/> |
| edgeR | 3.38.1 | https://bioconductor.org/packages/release/bioc/html/edgeR.html |
| extrafont | 0.18.0 | https://github.com/wch/extrafont |
| forcats | 0.5.2 | <https://forcats.tidyverse.org/> |
| ggplot2 | 3.4.0 | <https://ggplot2.tidyverse.org/> |
| ggrepel | 0.9.2 | <https://ggrepel.slowkow.com/> |
| ggVennDiagram | 1.2.2 | https://cran.r-project.org/web/packages/ggVennDiagram/index.html |
| Glimma | 2.6.0 | https://github.com/Shians/Glimma |
| limma | 3.52.2 | <https://bioconductor.org/packages/release/bioc/html/limma.html> |
| pheatmap | 1.0.12 | https://cran.r-project.org/web/packages/pheatmap/ |
| purrr | 0.3.5 | https://purrr.tidyverse.org/ |
| RColorBrewer | 1.1-3 | https://cran.r-project.org/web/packages/RColorBrewer/ |
| readr | 2.1.3 | https://readr.tidyverse.org/ |
| sf | 1.0-9 | https://cran.r-project.org/web/packages/sf/ |
| stringr | 1.4.1 | https://stringr.tidyverse.org/ |
| tibble | 3.1.8 | https://tibble.tidyverse.org/ |
| tidyr | 1.2.1 | https://tidyr.tidyverse.org/ |
| tidyverse | 1.3.2 | https://www.tidyverse.org/ |
